# Interspecies transmission from pigs to ferrets of antigenically distinct swine H1 influenza A viruses with loss in reactivity to human vaccine virus antisera as measures of relative zoonotic risk

**DOI:** 10.1101/2022.09.12.507661

**Authors:** J. Brian Kimble, Carine K. Souza, Tavis K. Anderson, Zebulun W. Arendsee, David E. Hufnagel, Katharine M. Young, Nicola S. Lewis, C. Todd Davis, Amy L. Vincent Baker

## Abstract

During the last decade, endemic swine H1 influenza A viruses (IAV) from six different genetic clades of the hemagglutinin gene caused zoonotic infections in humans. The majority of zoonotic events with swine IAV were restricted to a single case with no subsequent transmission. However, repeated introduction of human-seasonal H1N1, continual reassortment between endemic swine IAV, and subsequent drift in the swine host resulted in highly diverse swine IAV with human-origin genes that may become a risk to the human population. To prepare for the potential of a future swine-origin IAV pandemic in humans, public health laboratories selected candidate vaccine viruses (CVV) for use as vaccine seed strains. To assess the pandemic risk of contemporary US swine H1N1 or H1N2 strains, we quantified the genetic diversity of swine H1 HA genes, and identified representative strains from each circulating clade. We then characterized the representative swine IAV against human seasonal vaccine and CVV strains using ferret antisera in hemagglutination inhibition assays (HI). HI assays revealed that 1A.3.3.2 (pdm) and 1B.2.1 (delta-2) demonstrated strong cross reactivity to human seasonal vaccines or CVVs. However, swine IAV from three clades that represent more than 50% of the detected swine IAVs in the USA showed significant reduction in cross-reactivity compared to the closest CVV virus: 1A.1.1.3 (alpha-deletion), 1A.3.3.3-clade 3 (gamma), and 1B.2.2.1 (delta-1a). Representative viruses from these three clades were further characterized in a pig-to-ferret transmission model and shown to exhibit variable transmission efficiency. Our data prioritize specific genotypes of swine H1N1 and H1N2 to further investigate in the risk they pose to the human population.

**Importance:** Influenza A virus (IAV) is endemic in both humans and pigs and there is occasional bidirectional transmission of viruses. The process of interspecies transmission introduces novel viruses that increases the viral diversity in each host, impacting viral ecology and challenging control efforts through vaccine programs. Swine-origin IAVs have the potential to cause human pandemics, and pandemic preparation efforts include the identification and generation of candidate vaccine viruses (CVV) derived from epidemiologically relevant swine IAV surface proteins. The CVVs are derived from swine IAV detected and isolated in humans, and are updated infrequently; consequently the efficacy of these vaccines against contemporary swine IAV is unclear given rapid turnover and change of diversity. In this report we perform a risk assessment of contemporary swine H1 IAVs, determine whether current CVVs cross-react, and illustrate how swine-origin IAV replicate, transmit, and cause disease in a swine-to-ferret model system. In doing so, we identify the swine IAV that have lost cross-reactivity to current pandemic preparedness vaccines and demonstrate the utility of swine-to-ferret transmission experiments to further inform risk assessment.

## 1 Introduction

Influenza A viruses (IAV) infect a broad range of wild and domestic animal species and humans, and result in disease states ranging from asymptomatic to severe pneumonia and death. Wild waterfowl act as the natural reservoir for IAV, but various subtypes and lineages are endemic in domestic poultry, swine, and human populations. IAVs cause significant economic impact on swine productions systems (1–3). In the United States (US), three primary subtypes of IAV endemically circulate in swine populations with multiple hemagglutinin (HA) genetic clades present within each subtype (4). Within the H1 subtype, which accounted for approximately 68% of all IAVs isolated from US pigs in 2020, 8 HA phylogenetic clades and 9 neuraminidase (NA) clades were detected (4, 5). This considerable diversity is driven by a number of factors, including intra-species genetic evolution, reassortment, and the repeated introduction of human-origin IAV strains to pig populations (6–8). Additionally, a large diverse swine IAV population may have significant impact on human health, where swine-origin IAV may zoonotically transmit sporadically, termed “variant” in humans, or cause pandemics infecting millions of people, as seen during the 2009 H1N1 pandemic (H1N1pdm09) (9).

Swine H1 IAV in the US are classified by hemagglutinin (HA), and are either 1A lineage that evolved from the 1918 H1N1 pandemic, or 1B lineage that resulted from introduction and subsequent persistence of pre-2009 human seasonal H1N1 (10). The 1B lineage has 3 genetic clades that are currently circulating in the US: 1B.2.1, 1B.2.2.1, and 1B.2.2.2 (10). The 1A lineage viruses include 5 genetic clades that are currently circulating: 1A.1.1.3, 1A.2, 1A.2-3-like, 1A.3.3.2, and 1A.3.3.3 (10, 11). Within the 1A lineage is the H1N1pdm09, with the HA assigned the global nomenclature of clade 1A.3.3.2. This clade of viruses emerged in swine, zoonotically infected humans and has since become endemic, replacing the existing seasonal H1N1 in humans (12, 13). Thus, the swine-origin pandemic 1A.3.3.2 clade has gained sustained transmission, evolution, and adaptation in the human population (13). Over the years since 2009, the 1A.3.3.2 human viruses were repeatedly reintroduced into swine herds (12), and have increased diversification of other swine HA clades by constantly adding human-origin internal genes to endemic swine H1 via reassortment (14). Thus, reverse-zoonoses alters and enhances viral diversity in swine, and potentially impacts the likelihood of zoonotic infection through the pairing of human-origin genes to antigenically unique swine surface proteins.

Since 2009, efforts increased to prepare for the next potential IAV pandemic of swine origin. Applying the lessons learned from generating the H1N1pdm09 human vaccine, a selection of variant IAV from human zoonotic isolates have been used to create candidate vaccine viruses (CVV) (15, 16). If a swine-origin variant IAV emerged in the human population, a CVV could be used as seed stock to rapidly initiate vaccine production, provided there was antigenic cross-reactivity between the novel strain and the CVV and the variant IAV. Upon initial selection and generation, CVVs typically exhibit high cross reactivity to genetically similar viruses in swine (16). However, evolution of IAV in the swine host can result in antigenic change that will reduce the efficacy of CVVs. Further, of the eight swine H1 clades currently circulating in the US, only five have an available CVV, and there is limited understanding of how well those CVVs react with the diverse array of contemporary swine viruses.

In a previous study, we demonstrated that swine H1 lineage strains from 2012-2019 were significantly different from human seasonal vaccine strains and this antigenic dissimilarity increased over time as the viruses evolved in swine (11, 17). Pandemic preparedness CVV strains also demonstrated a loss in similarity with tested swine strains. Human sera revealed a range of responses to swine H1 IAV, including two lineages of viruses with little to no immunity, 1A.1.1.3 and 1B.2.1 (11). In this study, to further assess these swine H1, we identified contemporary, representative swine IAVs collected from 2019-2020. Selected viruses were tested against ferret antisera as a proxy for predicting the efficacy of available seasonal vaccines and CVV against current circulating swine IAVs. Of the tested strains, three swine H1 IAVs demonstrated reduced cross-reactivity to relevant CVVs and were derived from genetic clades that are frequently detected in surveillance. These strains were used in a pig-to-ferret transmission model to assess zoonotic transmission potential. This work uses *in silico, in vitro*, and *in vivo* approaches and identified gaps in current pandemic preparedness vaccine strategies by identifying three swine-origin H1 IAVs of zoonotic concern with a natural host species-based risk assessment.

## 2 Material and Methods

### 2.1 Genetic analysis and strain selection

Human IAV vaccine composition and pandemic preparedness CVV assessments occur biannually at the WHO Vaccine Composition Meeting. In these meetings, animal influenza activity data are presented with 6-month windows along with human seasonal influenza activity data. Consequently, we downloaded all available swine HA H1 sequences that were collected and/or deposited in GISAID between Jan 1, 2020 and June 30, 2020 (18). These sequences were aligned alongside CVV strains and human seasonal vaccine strains with mafft v7.453 (19), and each HA gene was classified to genetic clade within the octoFLU pipeline (20); if whole genome data were available for a strain, each gene was similarly classified to evolutionary lineage to determine genome constellation. Following classification, sequences were translated to amino acid, and a consensus HA1 for each identified clade was generated using flutile (https://github.com/flu-crew/flutile). A pairwise distance matrix was generated in Geneious Prime, and a wildtype field strain that was the best match to the HA1 clade consensus and that was available in the USDA IAV in swine virus repository was selected for additional characterization by hemagglutinin inhibition (HI) assay. Amino acid differences between CVVs or human seasonal vaccine strains and characterized swine IAV and clade consensus HA1s were generated using flutile (https://github.com/flu-crew/flutile). These data were visualized through the inference of a maximum-likelihood phylogeny for the HA nucleotide alignment using IQ-TREE v2 implementing automatic model selection (21). After selecting strains to represent contemporary swine H1 clades, selection criteria were expanded to include representative neuraminidase (NA) and internal gene constellations, the predominant evolutionary lineages were identified using octoFLUshow (5) and representative strains were tested against human seasonal vaccine and CVV ferret anti-sera.

### 2.2 Viruses and ferret antisera

Selected H1N1 and H1N2 isolates were obtained from the National Veterinary Services Laboratories (NVSL) through the U.S. Department of Agriculture (USDA) IAV swine surveillance system in conjunction with the USDA-National Animal Health Laboratory Network (NAHLN). Viruses used in this study were: 1A.1.1.3 A/swine/North Carolina/A02245416/2020 (NC/20) and A/swine/Texas/A02245420/2020 (TX/20); 1A.3.3.2 A/swine/Utah/A02432386/2019 (UT/19); 1A.3.3.3 A/swine/Minnesota/A02245409/2020 (MN/20); 1B.2.2.1 A/swine/Iowa/A02478968/2020 (IA/20); 1B.2.2.2 A/swine/Colorado/A02245414/2020; and 1B.2.1 A/swine/Illinois/A02139356/2018 (IL/18). Vaccine viruses were provided by the Centers for Disease Control and Prevention (CDC), Atlanta, Georgia, USA and included 1A.1.1.3. IDCDC-RG59 A/Ohio/24/2017-CVV (OH/24/17), 1A.3.3.3 A/Ohio/9/2015 (OH/15), 1A.3.3.2 A/Idaho/7/2018 (ID/18), 1B.2.2.1 A/Iowa/32/2016 (IA/16), 1B.2.1 A/Ohio/35/2017 (OH/35/17), and 1B.2.1 A/Michigan/383/2018 (MI/18). Viruses were grown in Madin-Darby canine kidney (MDCK) cells in Opti-MEM (Life Technologies, Waltham, MA) with 10% fetal calf serum and antibiotics/antimycotics supplemented with 1g/ml tosyl phenylalanyl chloromethyl ketone (TPCK)-trypsin (Worthington Biochemical Corp., Lakewood, NJ).

Ferret antisera produced against candidate virus vaccine (CVV) strains were kindly provided by CDC, Atlanta, Georgia, U.S. Antisera raised in ferrets against the following viruses were used: 1A.1.1.3 IDCDC-RG59 A/Ohio/24/2017-CVV, 1A.3.3.3 IDCDC-RG48 A/Ohio/9/2015-CVV, 1A.3.3.2 A/Idaho/7/2018, 1B.2.2.1 A/Iowa/32/2016, 1B.2.1 A/Ohio/35/2017, and 1B.2.1 A/Michigan/383/2018.

### 2.3 Hemagglutination Inhibition

Ferret antisera were heat inactivated at 56°C for 30 min then treated with a 20% Kaolin suspension (Sigma-Aldrich, St.Louis, MO) followed by adsorption with 0.75% guinea pig red blood cells (gpRBC) to remove nonspecific hemagglutination inhibitors as previously described (17). Treated ferret antisera were used in HI assay with gpRBCs. Briefly, 4 HAU of virus in 25ul was mixed with 25ul of two-fold serially diluted serum. After a 30-minute incubation at room temperature, 50ul of 0.75% gpRBCs were added and allowed to settle for 1 hour. Wells were observed for hemagglutination activity and the reciprocal of the highest serum dilution factor that prevented hemagglutination was recorded as the HI titer.

### 2.4 Swine-to-ferret transmission study design

Twenty 3-week-old piglets of mixed sex were obtained from an IAV- and porcine reproductive and respiratory syndrome virus-free herd. Prophylactic antibiotics (Excede; Zoetis, Florham Park, NJ) were administered upon arrival to prevent potential respiratory bacterial infections. Sixteen 4–6-month-old male and female ferrets were obtained from an influenza-free high health source. Animals were housed under BSL2 containment in compliance with the USDA-ARS NADC institutional animal care and use committee. Serum was collected from each pig and ferret and screened by a commercial enzyme-linked immunosorbent assay (ELISA) (MultiS ELISA; Idexx, Westbrook, ME) prior to experimental manipulations to confirm all animals were free of prior immunity and maternally acquired IAV specific antibodies. Pigs were divided randomly into groups of 5 and placed into separate containment rooms. Three groups received 2ml of IAV inoculum at 1 × 10^6^ TCID_50_ via intranasal administration. At two days post inoculation (dpi) four ferrets were placed in the room in separate, open-fronted isolators placed approximately 4 feet from pig decking (22). All animals received a subcutaneous radio frequency microchip (pigs: Deston Fearing, Dallas, TX; Ferrets: Biomedic Data Systems Inc., Seaford, DE) for identification and body temperature monitoring purposes. Body temperature and weight (ferrets only) were recorded from −3 to 14 dpi, with the readings recorded prior to exposure used for establishing a baseline reading. Ferrets were provided routine care and handled before pigs, with a change in outer gloves and decontamination of equipment with 70% ethanol between individual ferrets.

Three pigs from each experimental group were euthanized at 5dpi and necropsied to evaluate lung lesions and collect broncho-alveolar lavage fluid (BALF) (23). All pigs were nasal swabbed at 0, 1, 3, and 5 dpi as previously described. The remaining pigs were swabbed on 7 and 9 dpi and euthanized at 14dpi. Blood samples were collected prior to exposure and at necropsy for all pigs.

Contact ferrets were sampled by nasal wash collection at 0, 1, 3, 5, 7, 9, 11 and 12-days post contact (dpc) (22). BALF samples from ferrets were collected at necropsy (12 dpc) (22). Blood samples were collected prior to exposure and at 12 dpc to test for seroconversion by HI assays and (17, 24) by a commercial NP-ELISA (MultiS ELISA; Idexx, Westbrook, ME).

### 2.5 Virus replication and shedding

Nasal samples and BALF samples were titrated on MDCK cells to evaluate virus replication in the nose and lungs, as previously described (23). Inoculated monolayers were evaluated for cytopathic effect (CPE) between 48 and 72 h post-infection, and positive wells were identified by testing supernatant via hemagglutination assay with turkey RBC. A TCID_50_/ml titer was calculated for each sample using the method described by Reed and Muench (25).

### 2.6 Pathology examination

Swine lungs were evaluated for lesions at 5 dpi following standard protocols to assess pathogenesis in swine and potential for transmission to ferrets (23). The percentage of the lung affected with pneumonic consolidation typical of influenza virus in ferrets was visually estimated at 12 dpc to assess resolution of disease following transmission, following methods of scoring previously described (22). Tissue samples from the trachea and right middle or affected lung lobe were fixed in 10% buffered formalin for histopathologic examination. Tissues were processed by routine histopathologic procedures and slides stained with hematoxylin and eosin (H&E) or stained by immunohistochemistry.

### 2.7 Microbiological assays

Swine BALF samples were cultured for aerobic bacteria on blood agar and Casmin (NAD-enriched) plates to indicate the presence of concurrent bacterial pneumonia. To exclude other causes of pneumonia in pigs, PCR assays were conducted for porcine circovirus 2 (PCV2) (26), and for *Mycoplasma hyopneumoniae* and North American and European PRRSV (VetMax; Life Technologies, Carlsbad, CA) according to the manufacturer’s recommendations.

## 3 Results

### 3.1. Genetic and phylogenetic characterization of US swine H1 hemagglutinin

Between January 1 2020 and June 30 2020, 342 swine IAV isolates with an H1 HA were identified. These viruses represented 8 genetic clades across two evolutionary lineages: 1A.1.1.3 (n=36, 10.6%), 1A.2 (n=3, 0.9%), 1A.2-3-like (n=3, 0.9%), 1A.3.3.2 (n=53, 15.5%), 1A.3.3.3 (n=145, 42.4%), 1B.2.1 (n=80, 23.4%), 1B.2.2.1 (n=15, 4.4%), 1B.2.2.2 (n=7, 2%) (Figure 1). For each detected H1 clade, HA1 amino acid sequences were aligned, a consensus sequence generated, and a wildtype virus with highest HA1 similarity to consensus was selected to represent the clade (Table 1). The percent amino acid identity of selected strains ranged from 96.63-99.39% when compared to matching within-clade consensus and ranged from 90.83-98.16% when compared to within-clade CVVs (Table 1). The number of amino acid differences between the representative HA gene and the within-clade CVV or human seasonal vaccine ranged from 6 to 29 amino acid differences (Supplemental Tables 1-6).

**Figure 1.**
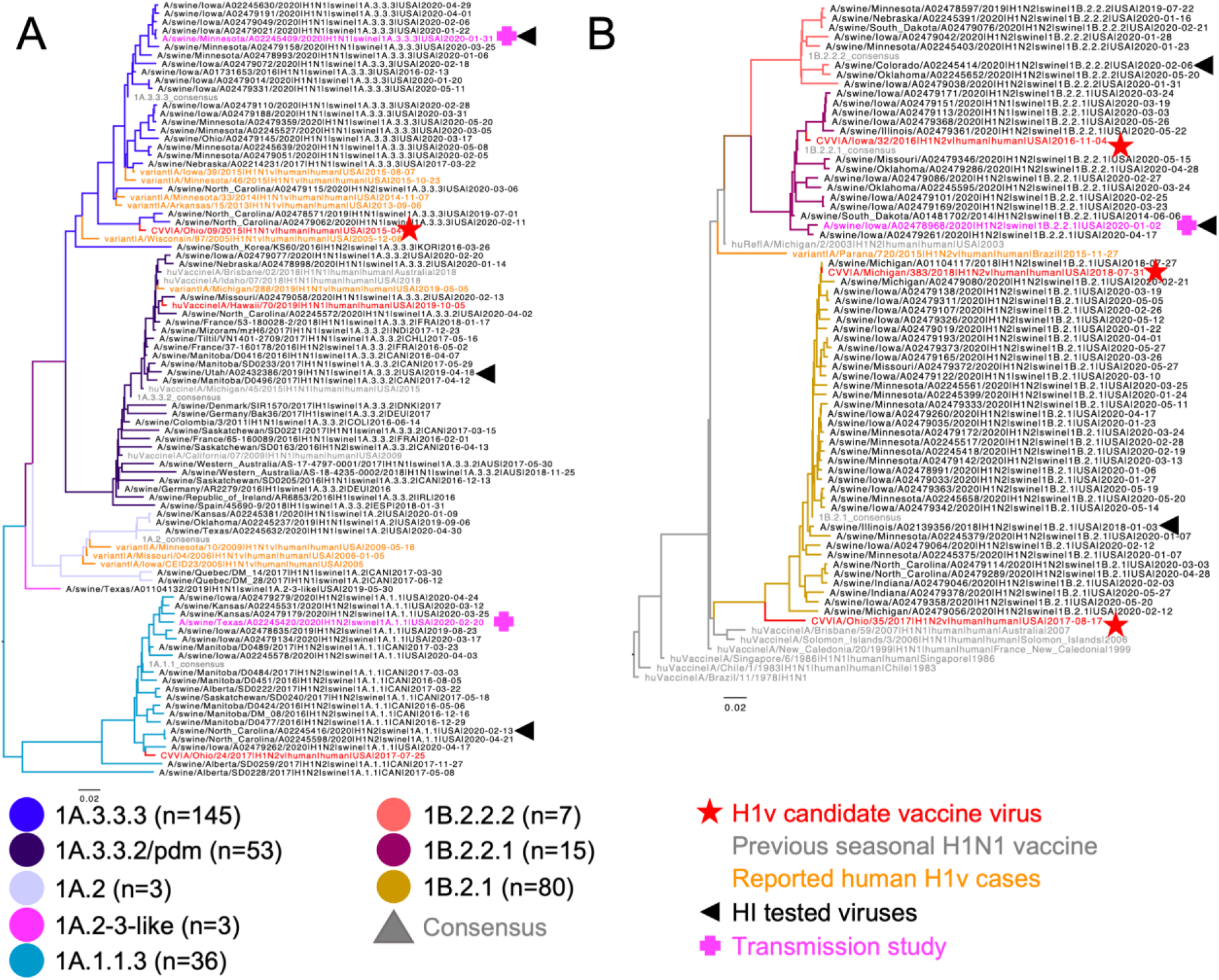
Phylogenetic relationships of North American swine H1 IAV. A representative random sample of (A) 1A classical swine lineage and (B) 1B human-like lineage swine HA genes from January 2020 through June 2020. Reference human HA genes, candidate vaccine viruses (CVV), and variant cases are indicated by branch color or shapes. Swine IAV strains tested in hemagglutination inhibition assays are marked by a black triangle and those used in transmission studies by a pink plus (+). The numbers in parentheses indicate number of each genetic clade detected during the sampling period.

**Table 1.**
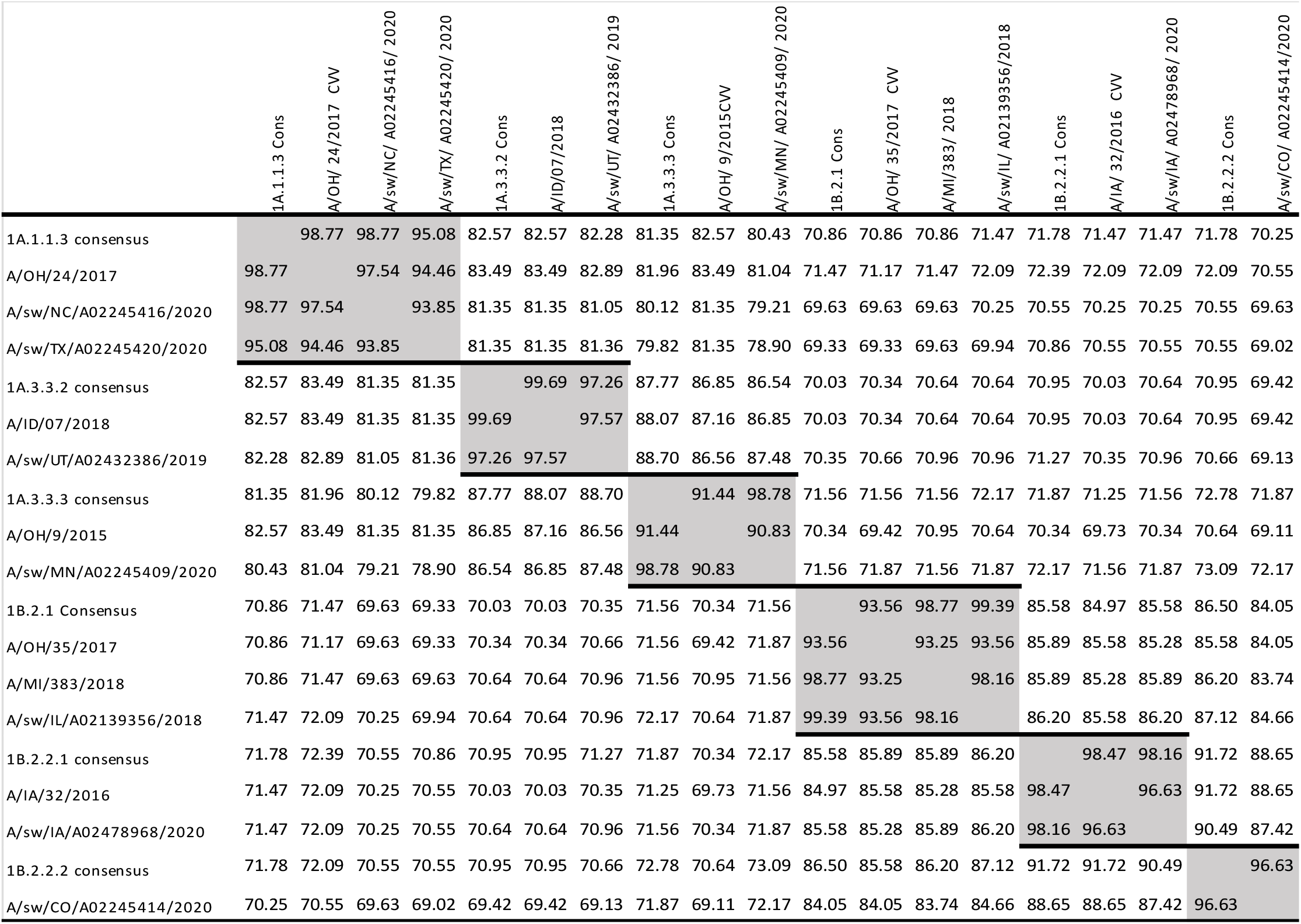
Pairwise amino acid sequence similarity of the HA1 domain from swine H1 clade consensus sequences to candidate vaccine viruses or seasonal vaccine virus and the swine hemagglutinin clade representative viruses used in this study. Within-clade comparisons are highlighted in grey.

### 3.2 Dominant U.S. swine H1 strains drifted from human seasonal H1 or CVV

In addition to current human seasonal vaccines, CVV strains were selected and generated by WHO collaborating centers to mitigate a future potential outbreak of swine IAV in humans. Ferret antisera generated against human vaccine and CVV strains and other variant IAV viruses were tested by HI to determine the relative cross-reactivity to contemporary swine viruses. Within the 1A lineage of HA genes, there were 2 CVVs and the human seasonal H1pdm vaccine strain that correspond to the 1A1.1, 1A.3.3.3 and 1A.3.3.2 clades, respectively (Table 2). The consensus 1A1.1 contemporary representative virus, NC/20, had an HI titer of 80 against CVV OH/24/17 antiserum compared to the homologous OH/24/17 titer of 1280, representing a 16-fold reduction in cross-reactivity. Similarly, the 1A.3.3.3 contemporary representative MN/20 virus displayed a 32-fold reduction in HI activity compared to homologous 1A.3.3.3 CVV OH/15. Conversely, the 1A.3.3.2 selected virus, UT/19, had had no loss in HI titer as compared to the homologous ID/18 (A/Brisbane/02/2018 (H1N1)-like) human strain.

**Table 2.** Antigenic cross-reactivity of 1A viruses and within-clade CVVs. Ferret antisera raised against CVV and vaccine viruses were tested for the ability to inhibit hemagglutination of contemporary swine viruses. Vaccine strains and homologous titers are bolded; grey highlighted cells indicate the within clade titer of contemporary swine strain.

| Strain | Lineage | IDCDC-RG59 A/Ohio/24/2017-CVV | IDCDC-RG48 A/Ohio/9/2015-CVV | A/Idaho/7/2018* |
| --- | --- | --- | --- | --- |
| **A/Ohio/24/2017** | **1A.1.1.3** | **1280** | <10 | 640 |
| A/swine/North Carolina/A02245416/2020 | 1A.1.1.3 | 80 | <10 | <10 |
| A/swine/Oklahoma/A02245237/2019 | 1A.2 | 20 | 320 | 640 |
| **A/Ohio/9/2015** | **1A.3.3.3** | <10 | **2560** | <10 |
| A/swine/Minnesota/102245409/2020 | 1A.3.3.3 | <10 | 80 | 40 |
| **A/Idaho/7/2018\*** | **1A.3.3.2** | <10 | <10 | **1280** |
| A/swine/Utah/A02432386/2019 | 1A.3.3.2 | 40 | 160 | 2560 |

\*A/Brisbane/02/2018 (H1N1)-like.

The 1B lineage of HA genes also had three antisera generated against variant viruses used to generate CVVs: 1B.2.2.1 IA/16 and 1B.2.1 OH/35/17 and MI/18. The 1B.2.2.1 virus, IA/20, had an 8-fold reduction in cross-reactivity compared to the 1B.2.2.1 CVV IA/16 virus. There is no clade-specific CVV for the 1B.2.2.2 clade of swine viruses and the representative virus, CO/20 had limited reactivity to all potential vaccine sera. Finally, the 1B.2.1 virus, IL/18, was antigenically very similar (2-fold or less reduction) to both 1B.2.1 CVV antisera.

### 3.3 Swine-to-ferret transmission

The antigenic data indicated the three clades of swine IAV with lowest reactivity to vaccine antisera: 1A.1.1.3, 1A.3.3.3, and 1B.2.2.1. These 3 clades accounted for 54.2% of H1 subtype IAV swine isolates during 2020 (5). Therefore, viruses from 1A.1.1.3, 1A.3.3.3, and 1B.2.2.1 clades were selected to test the zoonotic potential using a pig-to-ferret interspecies transmission model. While antigenicity is primarily driven by genetic factors within the HA1 domain of the HA gene, zoonosis is affected by viral factors attributed throughout the genome. To address this, we expanded our selection criteria to include NA and internal gene constellations (n = 225 H1N1 and H1N2 whole genome sequences collected in 2020: Supplemental Figure 1). During 2020, the 1B.2.2.1 HA gene was primarily paired with a N2-2002B gene with a TTTTPT internal gene constellation; the 1A.3.3.3 HA gene was primarily paired with a N1-Classical gene with a TTTPPT internal gene constellation. The 1A.3.3.3 (A/Minnesota/A02245409/2020) and 1B.2.2.1 (A/swine/Iowa/A02478968/2020) viruses selected for the antigenic characterization matched the predominant circulating NA and dominant internal gene constellations and thus remained unchanged. For the 1A.1.1.3 HA clade, the primary NA pairing in 2020 was a N2-2002A gene with detections of TTTTPT, TTTPPT, TTPTPT, and TTPPPT. The NC/20 virus used for HI assays did not match the predominant N2-NA gene, and had a TTTPPT internal gene constellation, and consequently, the strain selected for subsequent *in vivo* studies was A/swine/Texas/A02245420/2020 (TX/20) that had a N2-2002A gene with a TTTTPT internal gene constellation.

All three selected viruses had similar shedding patterns in pigs in terms of peak and duration (Figure 2). Evaluation of samples collected at the 5dpi necropsy revealed similar levels of macroscopic pathology between the 1A.1.1.3 (alpha) and the 1B.2.2.1 (delta-1a) groups at 2.4 and 2.3 percent of affected lung surface respectively. These two groups had a mean BALF titer of 3.6×10^5^ TCID_50_/ml and 5.9×10^6^ TCID_50_/ml respectively. The 1A.3.3.3 (gamma) virus had higher average percentage of lung lesions (8.1%) and higher BALF titers (1.8×10^7^ TCID_50_/ml) than the other groups. All 6 remaining pigs seroconverted at 14dpi with an average HI titer of 905, 640, and 80 in 1A.1.1.3, 1A.3.3.3, and 1B.2.2.1, respectively. These data demonstrated the propensity for the pigs to seed the room with aerosolized virus to expose the ferrets.

**Figure 2.**
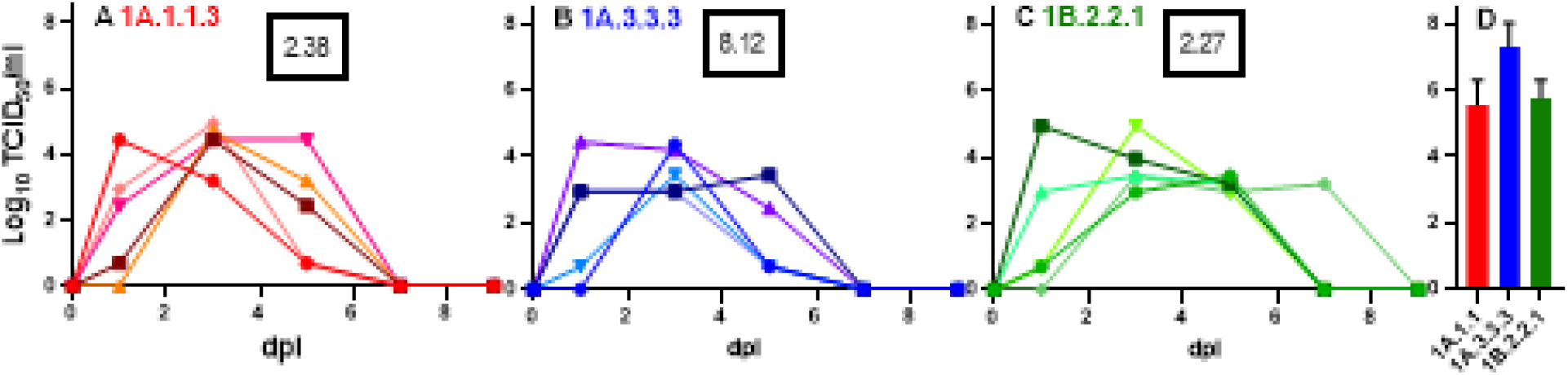
Shedding and replication of clade representative viruses in pigs. Individual pig nasal samples (A-C) and group mean bronchioalveolar lavage fluid (BALF) (D) viral load shown as TCID_50_ on MDCK cells. Results are reported as log10 TCID_50_/ml. Number in the black box indicates the average (n=3) percentage of lung surface with visible pneumonic lesions at the 5dpi necropsy.

The three viruses displayed different levels of transmissibility to contact ferrets (Figure 3). All four 1A.1.1.3 contact ferrets shed virus with an average of 2.3 positive sample days and an average peak titer of 7.9 ×10^5^ TCID_50_/ml in nasal washes. All four ferrets seroconverted with a geometric mean HI titer of 679 at 12dpc. The four 1A.3.3.3 contact ferrets all seroconverted as well, but with a lower geometric mean titer (231) and only two of the four ferrets had recoverable viral loads in the nasal washes with an average of 2.5 positive sample days and an average peak titer of 5.38×10^5^ in nasal washes. In contrast, only one of four 1B.2.2.1 contact ferrets seroconverted (HI=160) and had 2 nasal wash positive sample days with a peak titer of 8.9 ×10^4^ TCID_50_/ml. No ferret had recoverable viral loads in the BALF samples collected at 12dpc.

**Figure 3.**
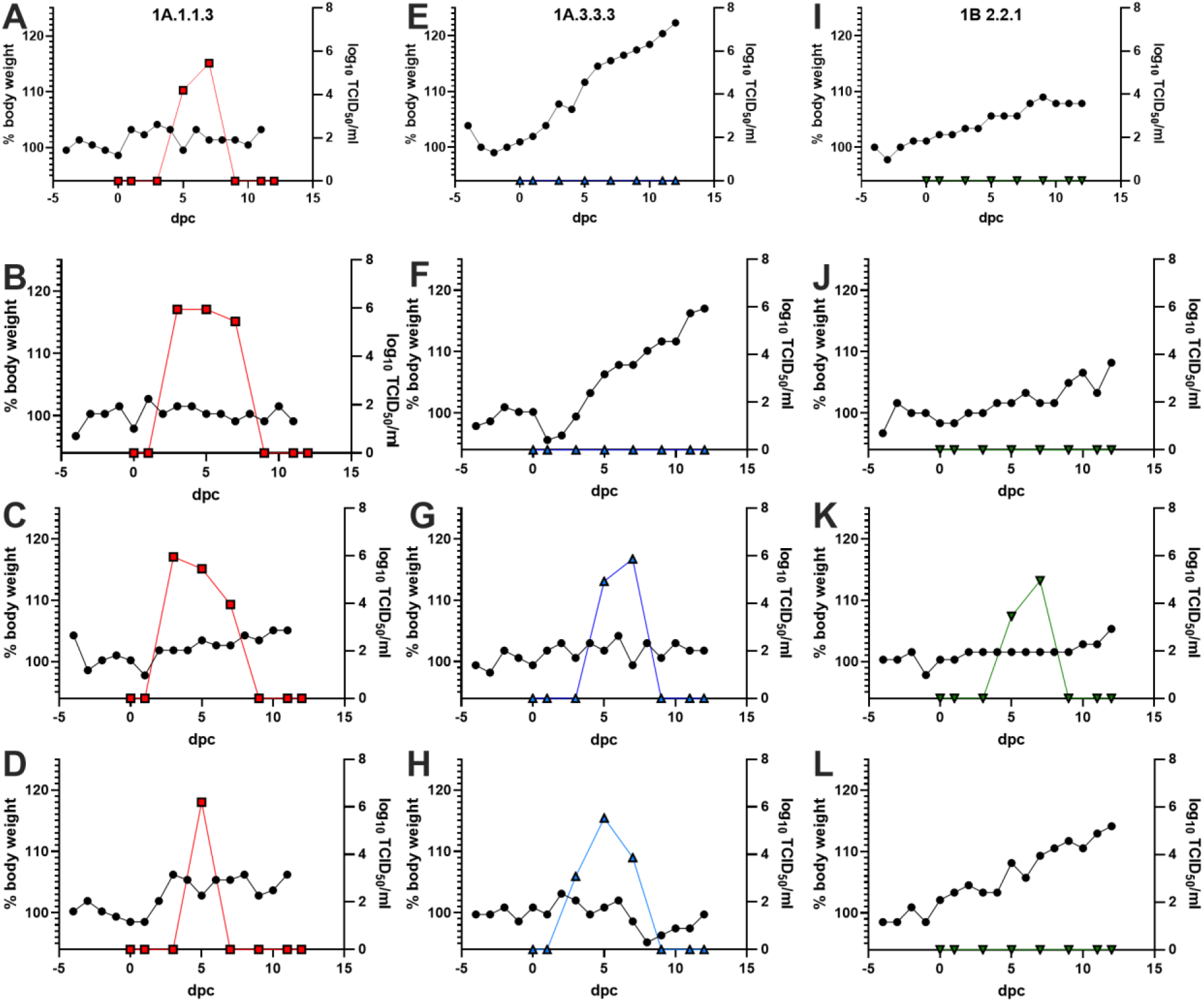
Transmission and replication of swine viruses to ferrets. Individual ferrets exposed to pigs infected with a 1A.1.1.3 (A-D, red squares), 1A.3.3.3 (E-H, blue up-triangles) or 1B.2.2.1(I-L, green down-triangles) had body weights recorded daily and converted to a percentage of the 3-day average body weight prior to exposure (left axis, black circles). Nasal washes were measured for viral shedding by TCID_50_ in MDCK cells and recorded as log10 TCID_50_/ml (right axis, color square).

Minimal signs of disease were observed in the ferrets. Daily temperature measurements revealed minimal elevation in temperatures and no differences among or between treatment groups. Body weight monitoring revealed if a ferret shed virus, regardless of group, it gained significantly less weight (3.5 ± 2.9%) compared to ferrets that did not shed virus (10.9 ± 6.5%) (Table 4), but there were no significant differences in change in body weight between virus groups. Necropsy on 12dpc revealed minimal gross pathology, with only two ferrets, one 1A.1.1.3 (4.6%) and one 1B.2.2.1 (3.5%), demonstrating visible lung lesions. No lesions were observed in any 1A.3.3.3 exposed ferrets at 12dpc (Table 4).

## Discussion

Animal origin influenza A viruses (IAV) from avian or swine are a documented source of human IAV zoonotic infections, epidemics, and pandemics. The four most recent IAV pandemics were all driven by either direct zoonosis or by reassortment and zoonosis (9, 27). Regional epidemics and smaller outbreaks were also initiated by the spillover of avian and swine viruses into human populations (https://www.cdc.gov/flu/weekly/fluviewinteractive.htm). In addition to the 2009 pandemic, swine-origin IAV are also responsible for annual human infections, termed variants, ranging from single cases up to outbreaks of several hundred. Much progress has been made in preparing for future animal origin IAV pandemics, with the most proactive efforts centered on the generation of CVV from human cases of IAV of animal origin, including variant viruses of swine origin. These stockpiled CVVs would be used as seed viruses for rapid vaccine generation should an antigenically similar animal origin virus initiate a human pandemic.

Swine IAV in the US is very diverse. In 2020 there were 14 antigenically distinct HA clades isolated from US swine herds, 8 of which were of the H1 subtype (4, 5). Contemporary clade consensus HA1 sequences can have as little as 70% amino acid similarity to divergent HA1 between swine H1 clades and within-clade HA1 sequences can be as much as 15% different (Table 1). This high level of within and between clade genetic diversity makes achieving and maintaining high levels of vaccine coverage difficult and necessitates the continued evaluation of CVV antisera reactivity against contemporary swine IAV isolates.

Of the 8 circulating swine H1 clades, five have existing seasonal or CVV vaccine options, including the 1A.3.3.2 component of human seasonal flu vaccine (15). Greater than 95% of 2020 US swine IAV isolates fall within those 5 human vaccine-covered clades (4, 5). Contemporary, clade-representative viruses of two of these clades showed high levels of cross-reactivity to existing vaccines, 1A.3.3.2 and 1B.2.1 (Table 2 & 3). Cross-species events involving human-to-swine infection of 1A.3.3.2 viruses in pigs are common in the US (6, 12). This continuous influx of human viruses makes it unsurprising that a representative swine 1A.3.3.2 virus had high levels of cross reactivity with a human 1A.3.3.2 vaccine. The 1B.2.1 A/Michigan/383/2018 is the most recently generated CVV. As such, it follows that the 1B.2.1 clade had not antigenically drifted and high levels of cross-reactivity were expected and observed. Three swine IAV clades have no within-clade vaccine or CVV available: 1A.2, 1A.4, and 1B.2.2.2. Additionally, these three clades had limited cross-reactivity to inter-clade CVVs. However, these three clades only represented 4.5% of 2020 US swine IAV isolates. This relative scarcity may minimize the opportunities for zoonotic transmissions and reduced the priority for assessing their pandemic risk posed to humans at this time. However, relative detection frequency of swine HA clades changes over time and these clades may need to be reassessed in the future given frequent interstate movement of pigs and viruses (28–31). Contemporary clade representative isolates from 1A.1.1.3 (16-fold reduction), 1A.3.3.3 (32-fold reduction) and 1B.2.2.1 (8-fold reduction) exhibited high levels of antigenic drift from relevant CVVs and these three H1 swine clades represented 54.2% of 2020 US swine IAV isolates (5). These clades have high frequency of detection in US swine herds and have reduced vaccine reactivity to human CVV, indicating a higher potential pandemic risk and requiring further examination of transmission risk factors. The 1A.1.1.3 and 1B.2.2.1 swine H1 clades also showed low detection by human population sera in a previous study (11). H3 clades represented 32% of all IAV in swine detected in 2020 (n=346 of 1093,(5)), but these H3 will be assessed in a separate publication.

**Table 3.** Antigenic cross-reactivity of 1B viruses and intra-clade CVVs. Ferret antisera raised against candidate vaccine viruses were tested for the ability to inhibit hemagglutinin of contemporary swine viruses. Vaccine strains and homologous titers are bolded; grey highlighted cells indicate the within-clade titer of contemporary swine strains.

| Strain | Lineage | A/Iowa/32/2016 | A/Ohio/5/2017 | A/Michigan/383/2018 |
| --- | --- | --- | --- | --- |
| **A/Iowa/32/2016** | 1B.2.2.1 | **640** | 20 | 10 |
| A/swine/Iowa/A02478968/2020 | 1B.2.2.1 | 80 | 10 | 40 |
| A/swine/Colorado/A02245414/2020 | 1B.2.2.2 | 40 | 10 | 10 |
| **A/Ohio/35/2017** | 1B.2.1 | 80 | **640** | 40 |
| **A/Michigan/383/2018** | 1B.2.1 | 40 | 160 | **1280** |
| A/swine/Illinois/A02139356/2018 | 1B.2.1 | 20 | 320 | 1280 |

To address zoonotic potential, these viruses were used in a swine-to-ferret interspecies transmission study. While all three viruses exhibited some level of interspecies transmission, they did so with varying efficiency (Figure 3). The 1A.1.1.3 virus had complete transmission from pigs to ferrets indicated by all ferrets shedding virus and seroconverting. All four of the 1A.3.3.3 exposed ferrets also seroconverted, albeit with a lower average HI titer compared to the 1A.1.1.3, but only two of the four ferrets shed virus. Finally, one 1B.2.2.1 ferret seroconverted and shed virus while the other three remained naïve, indicating a reduced propensity for interspecies transmission. The infected ferrets displayed signs of disease measured as a cessation of weight gain compared to noninfected ferrets, but no other overt signs and postmortem evaluation of the lungs revealed minimal pathological damage at 12pc (Table 4). Since this study was focused on transmission rather than pathogenesis in ferrets, further work to determine lung pathology during the active infection phase would be necessary. Nasal titers over the time course and lung titers on 5 dpi in pigs were similar for all three viruses, indicating that infection and replication kinetics in pigs did not affect transmission to ferrets. The virus and host factors contributing to the lower nasal shedding of the 1A.3.3.3 and the lower transmission of the 1B.2.2.1 swine strains in the contact ferrets are currently unknown, but potentially associated with the diverse gene segment combinations and evolutionary origins of the three viruses.

**Table 4.** Cumulative clinical, viral, and serological measures of ferret infection.

| 1 | Ferret # | Change in<br>bodyweight<br>(%) from 0-12 DPC | DPC with<br>nasal<br>shedding | Peak nasal<br>titer* | 12 DPC HI<br>titer |
| --- | --- | --- | --- | --- | --- |
| 1A.1.1.3 | 1 | 4.61 | 5,7 | 5.45 | 160 |
|  | 2 | 1.19 | 3,5,7 | 5.94 | 1280 |
|  | 3 | 4.89 | 3,5,7 | 5.94 | 1280 |
|  | 4 | 7.71 | 5 | 6.2 | 1280 |
| 1A.3.3.3 | 5 | 21.36 | none | none | 80 |
|  | 6 | 16.83 | none | none | 160 |
|  | 7 | 2.40 | 5,7 | 5.87 | 320 |
|  | 8 | -1.13 | 3,5,7 | 5.53 | 1280 |
| 1B.2.2.1 | 9 | 6.74 | none | none | <10 |
|  | 10 | 9.84 | none | none | <10 |
|  | 11 | 5.02 | 5,7 | 4.95 | 160 |
|  | 12 | 12.01 | none | none | <10 |
| No virus | 13 | 7.25 | none | none | <10 |
|  | 14 | 2.35 | none | none | <10 |

\*Log<sub>10</sub> TCID<sub>50</sub>/ml

Results of this study indicate that swine IAV from the US may escape vaccine immunity from CVV or seasonal vaccines as they continue to circulate and evolve in the swine population. Three H1 clades demonstrated antigenic drift away from available CVV antisera. Additionally, contemporary clade representatives showed the ability to transmit from pigs to ferrets, a gold standard for human influenza transmissibility. These data highlight the increased risk to human populations posed by H1 clades of swine IAV, particularly the 1A.1.1.3. Since the conclusion of these experiments in July 2020, there have been 15 H1 variant cases in North America where the HA clade could be determined; an additional 3 variants had insufficient data to identify the HA clade. Of these variant IAVs, 2 came from the 1A.1.1.3 clade, 4 were derived from the 1A.3.3.3 clade, 5 were from the 1A.3.3.2 clade, and 4 were from the 1B.2.1 clade. These data highlight the utility of swine-to-ferret transmission studies as a pandemic risk assessment tool and identifies the gaps in CVV coverage of US H1 swine IAV. The results stress the need to continually assess the intra-clade cross-reactivity of existing CVVs to identify and develop more contemporarily relevant pandemic preparedness strains.

## Acknowledgments

We gratefully acknowledge pork producers, swine veterinarians, and laboratories for participating in the USDA Influenza A Virus in Swine Surveillance System and publicly sharing sequences. We also gratefully acknowledge all data contributors, i.e., the Authors and their Originating laboratories responsible for obtaining the specimens, and their Submitting laboratories for generating the genetic sequence and metadata and sharing via the GISAID Initiative, on which components of this research is based. This work was supported in part by: the U.S. Department of Agriculture (USDA) Agricultural Research Service [ARS project number 5030-32000-231-000-D]; the National Institute of Allergy and Infectious Diseases, National Institutes of Health, Department of Health and Human Services [contract numbers HHSN272201400008C and 75N93021C00015]; the USDA Agricultural Research Service Research Participation Program of the Oak Ridge Institute for Science and Education (ORISE) through an interagency agreement between the U.S. Department of Energy (DOE) and USDA Agricultural Research Service [contract number DE-AC05-06OR23100]; and the SCINet project of the USDA Agricultural Research Service [ARS project number 0500-00093-001-00-D]. The funders had no role in study design, data collection and interpretation, or the decision to submit the work for publication. Mention of trade names or commercial products in this article is solely for the purpose of providing specific information and does not imply recommendation or endorsement by the USDA, CDC, DOE, or ORISE. USDA and CDC are equal opportunity providers and employers. The findings and conclusions in this report are those of the authors and do not necessarily represent the views of the Centers for Disease Control and Prevention or the Agency for Toxic Substances and Disease Registry.

