## Supplemental material for "Interspecies transmission from pigs to ferrets of antigenically distinct swine H1 influenza A viruses with loss in reactivity to human vaccine virus antisera as measures of relative zoonotic risk"

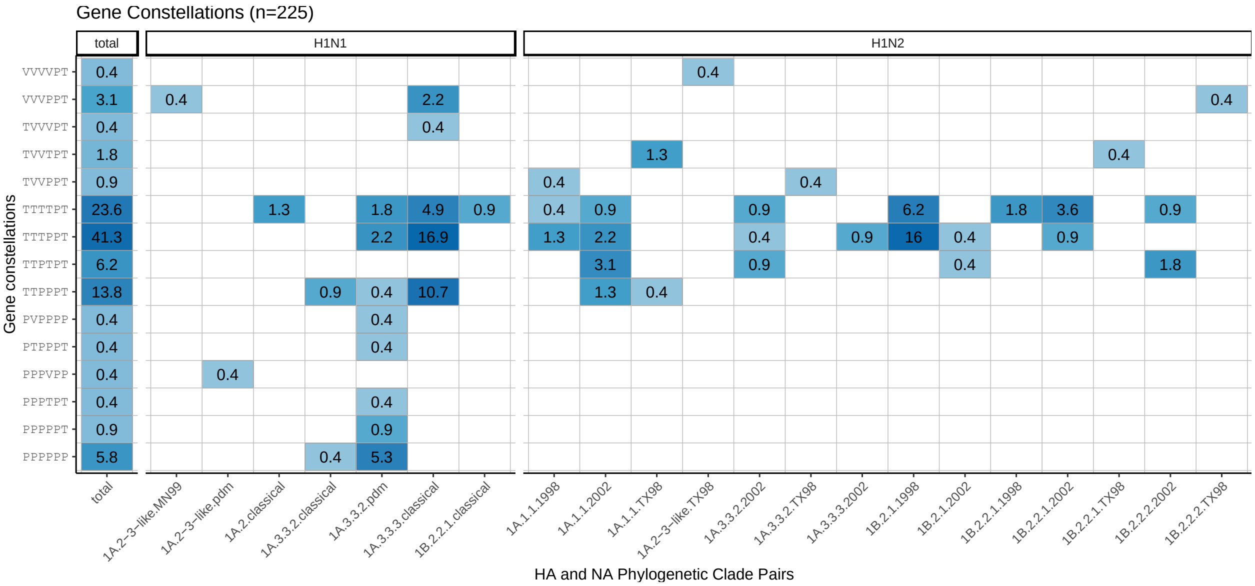


**Supplemental Figure 1.** Detection proportions of H1N1 and H1N2 influenza A virus in swine collected in 2020 in the USDA influenza A virus in swine surveillance system. The x-axis reflects the paired genetic clade of hemagglutinin (HA) and neuraminidase (NA), and the y-axis reflects the evolutionary lineage (triple-reassortant, T; H1N1pdm09, P; live attenuated vaccine virus associated, V) of the internal genes in the order of PB2-PB1-PA-NP-M-NS.

**Supplemental Table 1.** Listing of amino acid differences between 1A.1.1.3 clade consensus, the within-clade CVV A/Ohio/24/2017, and the clade representative viruses A/swine/North Caroline/A02245416/2020 and A/swine/Texas/A02245420/2020.

| 1A.1.1.3 | H1 Numbering | AA# | **1A.1.1.3 Consensus** | A/Ohio/24/2017 CVV | A/swine/NC/A02245416/2020 | A/swine/Tx/A02245420/2020 |
|---|---|---|---|---|---|---|
| | 14 | 14 | **D** | | | E |
| | 48 | 48 | **A** | | | P |
| | 68 | 68 | **E** | | G | |
| | 119 | 119 | **M** | I | | I |
| | 126 | 126 | **Y** | H | | |
| | 132 | 130 | **V** | | | E |
| | 138 | 136 | **D** | | | Y |
| | 141 | 139 | **A** | | | R |
| | 142 | 140 | **S** | | | G |
| | 149 | 147 | **M** | I | | |
| | 156 | 154 | **D** | N | | |
| | 168 | 166 | **N** | | | D |
| | 170 | 168 | **R** | | | G |
| | 176 | 174 | **L** | | I | |
| | 209 | 207 | **E** | | | K |
| | 216 | 214 | **T** | | | I |
| | 224 | 222 | **T** | | | A |
| | 232 | 230 | **T** | | | A |
| | 235 | 233 | **E** | | K | |
| | 252 | 250 | **K** | | | R |
| | 264 | 262 | **G** | | S | |
| | 310 | 308 | **T** | | | R |
| | 311 | 309 | **K** | | | R |
| Differences from consensus | | | **-** | **4** | **4** | **15** |
| Differences from CVV | | | **4** | **-** | **8** | **16** |

**Supplemental Table 2.** Listing of amino acid differences between 1A.3.3.2 clade consensus, the within-clade human seasonal vaccine A/Idaho/07/2018, and the clade representative A/swine/Utah/A02432386/2019.

| 1A.3.3.2 | H1 numbering | **1A.3.3.2 Consensus** | A/Idaho/07/2018 | A/swine/UT/A02432386/2019 |
|---|---|---|---|---|
| | 47 | **V** | | I |
| | 74 | **R** | | S |
| | 164 | **T** | | S |
| | 183 | **P** | | S |
| | 205 | **R** | | K |
| | 216 | **T** | | S |
| | 260 | **D** | N | N |
| | 295 | **I** | | I |
| Difference from consensus | | **-** | **1** | **8** |
| Difference from vaccine | | 1 | **-** | **7** |

**Supplemental Table 3.** Listing of amino acid differences between 1A.3.3.3 clade consensus, the within-clade CVV A/Ohio/09/2015, and the clade representative virus A/swine/Minnesota/A02245409/2020.

| 1A.3.3.3 | H1 numbering | **1A.3.3.3 Consensus** | A/Ohio/09/2015 CVV | A/swine/MN/A02245409/2020 |
|---|---|---|---|---|
| | 2 | **T** | K | |
| | 3 | **L** | I | |
| | 36 | **R** | K | |
| | 71 | **S** | A | |
| | 83 | **S** | | P |
| | 84 | **N** | S | |
| | 86 | **E** | N | |
| | 113 | **R** | K | |
| | 120 | **T** | | A |
| | 127 | **D** | E | |
| | 137 | **P** | | S |
| | 146 | **R** | K | |
| | 149 | **V** | I | |
| | 153 | **Q** | K | |
| | 155 | **G** | E | |
| | 161 | **V** | I | |
| | 163 | **K** | I | |
| | 166 | **I** | T | |
| | 169 | **K** | R | R |
| | 170 | **E** | G | |
| | 183 | **S** | P | |
| | 186 | **A** | T | |
| | 192 | **K** | Q | |
| | 196 | **D** | N | |
| | 197 | **A** | S | |
| | 205 | **K** | R | |

| | | | |
|---|---|---|---|
| 221 | **D** | G | |
| 250 | **A** | V | |
| 269 | **D** | E | |
| 271 | **S** | P | |
| Differences from consensus | | **27** | **4** |
| Differences from CVV | **27** | | **29** |

**Supplemental Table 4.** Listing of amino acid differences between 1B.2.1 clade consensus, the within-clade CVVs A/Ohio/35/2017 and A/Michigan/383/2018, and the clade representative virus A/swine/Illinois/A02139356/2018.

| 1B.2.1 | H1 numbering | **1B.2.1 Consensus** | A/Ohio/35/2017 CVV | A/Michigan/383/2018 CVV | A/swine/IL/A02139356/2018 |
|---|---|---|---|---|---|
| | 36 | **S** | T | | |
| | 71 | **T** | I | N | |
| | 72 | **S** | P | | |
| | 83 | **S** | P | | |
| | 96 | **E** | T | | |
| | 113 | **K** | R | | |
| | 120 | **K** | D | | |
| | 121 | **S** | G | | |
| | 129 | **T** | S | | |
| | 156 | **N** | G | | |
| | 169 | **E** | K | | K |
| | 170 | **E** | | G | |
| | 173 | **I** | V | V | |
| | 185 | **M** | I | | |
| | 189 | **R** | K | | |
| | 208 | **R** | K | | |
| | 209 | **R** | K | | |
| | 216 | **K** | R | | |
| | 259 | **K** | | | R |
| | 260 | **G** | | S | |
| | 261 | **F** | S | | |
| | 277 | **T** | A | | |
| | 289 | **S** | D | | |
| | 310 | **A** | T | | |

| | | | | |
|---|---|---|---|---|
| Differences from consensus | **-** | **21** | **4** | **2** |
| Difference from OH/35 CVV | **21** | **-** | **22** | **21** |
| Difference from MI/383 CVV | **4** | **22** | **-** | **6** |

**Supplemental Table 5.** Listing of amino acid differences between 1B.2.2.1 clade consensus, the within-clade CVVs A/Iowa/32/2016 and the clade representative virus A/swine/Iowa/A02478968/2020.

| 1B.2.2.1 | H1 Numbering | **1B.2.2.1 consensus** | A/Iowa/32/2016 CVV | A/swine/IA/A02478968/2020 |
|---|---|---|---|---|
| | 19 | **V** | L | |
| | 74 | **K** | | E |
| | 84 | **N** | | D |
| | 96 | **A** | T | |
| | 132 | **V** | K | |
| | 141 | **K** | E | |
| | 166 | **K** | | E |
| | 168 | **D** | E | |
| | 237 | **G** | | R |
| | 273 | **D** | | N |
| | 277 | **A** | | T |
| Differences from consensus | | **-** | **5** | **6** |
| Differences from CVV | | **5** | **-** | **11** |

**Supplemental Table 6.** Listing of amino acid differences between 1B.2.2.2 clade consensus and the clade representative virus A/swine/Colorado/A02245414/2020.

| 1B.2.2.2 | H1 Numbering | **1B.2.2.2 Consensus** | A/swine/CO/A02245414/2020 |
|---|---|---|---|
| | 36 | **S** | N |
| | 50 | **L** | I |
| | 72 | **S** | P |
| | 85 | **S** | P |
| | 176 | **L** | I |
| | 208 | **G** | E |
| | 228 | **N** | K |
| | 236 | **P** | A |
| | 249 | **I** | V |
| | 271 | **P** | S |
| | 274 | **E** | K |
| Differences from Consensus | | **-** | **11** |
